# Generation of Pathogenic *TPP1* Mutations in Human Stem Cells as a Model for CLN2 Disease

**DOI:** 10.1101/2021.01.05.425495

**Authors:** Li Ma, Adriana Prada, Michael Schmidt, Eric M. Morrow

**Affiliations:** Department of Molecular Biology, Cell Biology and Biochemistry, Brown University, Providence, Rhode Island 02912, USA; Center for Translational Neuroscience, Carney Institute for Brain Science and Brown Institute for Translational Science, Brown University, Providence, Rhode Island 02912, USA; Hassenfeld Child Health Innovation Institute, Brown University, Providence, Rhode Island 02912, USA

**Author notes:** To whom correspondence should be addressed: Eric M. Morrow, M.D. Ph.D., Brown University, Laboratories for Molecular Medicine 70 Ship Street, Providence, Rhode Island 02912 USA.

**Keywords:** CLN2, TPP1, CRISPR/Cas9, Genome editing, Embryonic stem cells, Disease model

## Abstract

Neuronal ceroid lipofuscinosis type 2 (CLN2 disease) is an autosomal recessive neurodegenerative disorder generally with onset at 2 to 4 years of age and characterized by seizures, loss of vision, progressive motor and mental decline, and premature death. CLN2 disease is caused by loss-of-function mutations in the tripeptidyl peptidase 1 (*TPP1*) gene leading to deficiency in TPP1 enzyme activity. Approximately 60% of patients have one of two pathogenic variants (c.509-1G>C or c.622C>T [p.(Arg208*)]). In order to generate a human stem cell model of CLN2 disease, we used CRISPR/Cas9-mediated knock-in technology to introduce these mutations in a homozygous state into H9 human embryonic stem cells. Heterozygous lines of the c.622C>T (p.(Arg208*)) mutation were also generated, which included a heterozygous mutant with a wild-type allele and different compound heterozygous coding mutants resulting from indels on one allele. We describe the methodology that led to the generation of the lines and provide data on the initial validation and characterization of these CLN2 disease models. Notably, both mutant lines (c.509-1G>C and c.622C>T [p.(Arg208*)]) in the homozygous state were shown to have reduced or absent protein, respectively, and deficiency of TPP1 enzyme activity. These models, which we are making available for wide-spread sharing, will be useful for future studies of molecular and cellular mechanisms underlying CLN2 disease and for therapeutic development.

## 1. Introduction

The neuronal ceroid lipofuscinoses (NCLs)^1^, also known as Batten disease, are a group of neurodegenerative lysosomal storage disorders characterized by progressive intellectual and motor deterioration, visual failure, seizures, cerebellar atrophy, and premature death (Jalanko and Braulke, 2009; Kohlschutter and Schulz, 2009; Mole et al., 2018; Mole and Williams, 2001 [updated 2013]). Outside of one adult-onset type, NCLs are inherited in an autosomal recessive manner. The neuropathological features include aberrant accumulation of autofluorescent lipopigments in lysosomes. To date, mutations in 13 human genes have been linked with NCL disorders, which form the basis for classification as to NCL type (UCL, 2020).

CLN2 disease is caused by pathogenic variants in the tripeptidyl peptidase 1 (*TPP1*) gene leading to loss-of-function and deficiency in activity of the soluble lysosomal enzyme TPP1 (Junaid et al., 1999). Similar to other NCLs, it is characterized by an aberrant accumulation of autofluorescent lipopigments in lysosomal storage bodies in neurons, as well as in other cells, of patients (Sleat et al., 1997). Disease-causing variants appear along the whole length of the *TPP1* gene (Gardner et al., 2019). In CLN2 disease, approximately, 60% of patients have one of two pathogenic variants (c.509-1G>C and c.622C>T [p.(Arg208*)]). These mutations may be found in a homozygous state but are also commonly present in a compound heterozygous state (Gardner et al., 2019; Lojewski et al., 2014).

Lojewski et al. (2014) generated induced pluripotent stem cell (iPSC) models of CLN2 disease by reprogramming patient fibroblasts with compound heterozygous mutations in *TPP1*: c.[509-1G>C];[622C>T] (p.[(Arg208*)]). Similarly, Sima et al. (2018) generated iPSCs from two different patients with the following compound heterozygous mutations: c.[379C>T];[622C>T] (p.[(Arg127*)];[(Arg208*)] and c.[380G>A];[509-1G>C] (p.[(Arg127Gln]). Although these models are undoubtably of important utility in the CLN2 disease field, they present limitations. First, the context of compound heterozygous mutations does not easily permit mechanistic studies of specific individual mutations. Second and importantly, neither of these models to date have an associated isogenic control line, and thus, all studies were done using comparison to a genetically non-identical and unrelated control line.

In this study, we generated homozygous *TPP1* mutations for each of the two most common pathogenic variants, c.509-1G>C and c.622C>T (p.(Arg208*)), in H9 human embryonic stem cells (hESCs) using clustered regularly interspaced short palindromic repeats (CRISPR)/CRISPR-associated protein 9 (Cas9) genome editing. In generating these homozygous lines, heterozygous lines of the c.622C>T (p.(Arg208*)) mutation were also generated, which included a heterozygous mutant with a wild-type allele and different compound heterozygous mutants resulting from indels on one allele. We validated these cell lines and performed an initial characterization of TPP1 enzyme activity and cellular phenotypes. An advantage of our CLN2 disease models over prior work is that we have established each mutation in the homozygous state, which permits mechanistic and translational studies in the context of these specific individual mutations. Further, our models include the isogenic parental H9 hESC line as a control, which permits more direct assessment of effects resulting from the primary mutation.

## 2. Materials and methods

### 2.1. Genome editing of TPP1 in the H9 hESC line and Sanger sequencing

CRISPR/Cas9 homology directed repair knock-in technology was used for generating mutations at the *TPP1* locus in the H9 hESC line (WiCell #WA09). The RNA guide sequences were cloned into PX459 V2.0 plasmid (Addgene Plasmid #62988) digested with BbsI to allow for ligation of the guide RNA (gRNA) insert. Single-stranded oligodeoxyribonucleotide (ssODN) consisting of 100-nucleotide ultramers (Integrated DNA Technologies) was used as template to repair the double-stranded break. Detailed gRNA and ssODN sequences are summarized in Supplementary Table 1.

H9 hESCs were treated with 10 µM Rho-associated protein kinase inhibitor (ROCKi, Tocris Bioscience #Y-27632) for 24 h before they were dissociated to single cells and pelleted. Two µg of CRISPR plasmid and 5 µl of 100 mM ssODN were nucleofected into cells using Lonza 4D-Nucleofection Primary P3 Kit with program CB-150. The following day, puromycin at 0.5 µg/mL final concentration and ROCKi at 10 µM final concentration were added to fresh media, and cells continued to be cultured for 48 h. ROCKi was kept on the cells until small colonies were observed, which were clonal given their derivation from single cells. After this point, cells continued to be cultured with fresh media changes daily. After 15 days, colonies were isolated into 24-well plates. Four days after plating in the 24-well plates, a few colonies from each well were isolated and DNA was extracted using the HotSHOT method. PCR was performed across the targeted region using Herculase II (Agilent).

For the c.509-1G>C mutation, the point mutation from G to C created a BsrI restriction site. The PCR products were digested with BsrI, and complete digestion clones were further sequenced by Sanger sequencing to confirm the edit. Clones in which both alleles had been correctly targeted for the c.509-1G>C mutation (i.e., homozygous mutant clones) were identified in the first round of screening. For the c.622C>T mutation, to prevent the re-cutting of the genome, a silent mutation (c.630C>T [p.(Asn210Asn)]) was designed into the 3’ end of the mutation site in the ssODN. In designing the silent mutation, an MluCI restriction site was created in order to allow for screening for homologous recombination. The PCR products were digested with MluCI, and clones showing at least some level of digestion were further analyzed by Sanger sequencing to determine the edit. For this mutation, in the first round of screening, five clones were identified in which one allele was correctly targeted for the c.622C>T;630C>T mutations, whereas the other allele had an indel (four clones) or was wild-type (one clone). The latter clone, with one wild-type allele, was retargeted to generate homozygous mutant clones of the c.622C>T;630C>T mutations. Again, the PCR products were digested with MluCI, and complete digestion clones were further sequenced by Sanger sequencing to confirm the edit. Clones in which both alleles had been correctly targeted for the c.622C>T;630C>T mutations (i.e., homozygous mutant clones) were identified in the second round of screening. Detailed information concerning primers used for sequencing can be found in Supplementary Table 2. *TPP1*-mutant hESC lines containing homozygous versions of either the intronic splice acceptor mutation (c.509-1G>C) or the nonsense C>T mutation (c.622C>T;630C>T [p.(Arg208*;Asn210Asn)]) generated as part of this study have been deposited with the WiCell biorepository for distribution. The deposited cell lines have also been registered with hPSCreg (https://hpscreg.eu/), with identifiers as follows:

*TPP1* c.509-1G>C, clone #23; hPSCreg name: EMe-TPint5GC23

*TPP1* c.509-1G>C, clone #42; hPSCreg name: EMe-TPint5GC42

*TPP1* c.622C>T;630C>T (p.(Arg208*;Asn210Asn)), clone #12; hPSCreg name: EMe-TPR208X12

*TPP1* c.622C>T;630C>T (p.(Arg208*;Asn210Asn)), clone #25; hPSCreg name: EMe-TPR208X25

### 2.2. Maintenance of hESCs

H9 hESC lines, control and genome-edited, were cultured on Matrigel-coated plates (Corning #354277) under feeder-free conditions and maintained in mTeSR1 plus media (StemCell Technologies #85850). The medium was changed approximately daily. Enzyme-free passaging was performed using ReLeSR (StemCell Technologies #05872) according to the manufacturer’s instructions. Cell cultures were maintained at 37 °C in a humidified atmosphere of 95% air and 5% CO_2_.

### 2.3. Imaging of hESCs

Cultured H9 hESC lines were imaged using a Nikon Eclipse TS100 microscope equipped with a Q Imaging QIClick 1.4MP CCD monochrome microscope camera. Images were captured of cells being visualized using either a 4X or a 10X objective.

### 2.4. Western blotting

Cells for Western blotting (H9 hESCs and HEK293 cells) were harvested and then lysed in Flag lysis buffer (50 mM Tris-HCl, pH 7.8, 137 mM NaCl, 1 mM NaF, 1 mM NaVO_3_, 1% Triton X-100, 0.2% Sarkosyl, 1 mM dithiothreitol, and 10% glycerol) supplemented with protease inhibitor cocktail and phosphatase inhibitor for 30 min on ice. Cell lysates were generated by centrifugation at 13,200 rpm for 15 min at 4 °C. Protein concentration was measured by BCA assay using the Pierce BCA Kit (Thermo Fisher #23225). Samples containing 20 µg of protein were then boiled in sample buffer at 95 °C for 5 min before loading onto a 4-12% SDS-PAGE gel (Novex #NP0321Box). Following separation of proteins by electrophoresis, the gel was transferred to a nitrocellulose membrane (Novex #LC2000), and Western blot was performed using standard procedures (Ouyang et al., 2019; Xu et al., 2017). The Western blot was analyzed with the Li-CoR Odyssey Imaging System. Proteins were detected using mouse anti-TPP1 antibody (1:1000 working dilution; Abcam #ab54685) and mouse anti-α-Tubulin (1:5000 working dilution; Sigma #T6074). Bands were quantified by densitometry using ImageJ software (National Institutes of Health, https://imagej.nih.gov/ij/; three measurements per band), and the average measurement for a TPP1 band was normalized to the respective average measurement for an α-Tubulin band.

### 2.4. TPP1 enzyme activity assay

TPP1 enzyme activity was measured fluorometrically based on fluorescence of the chromophore 7-amino-4-methylcoumarin (AMC) (Sigma #A9891) as described previously (Lojewski et al., 2014), but with some modifications. Fifteen µg of cell lysate from each hESC line was incubated in 150 µl acetate buffer (pH 4.0) containing a final concentration of 62.5 µM Ala-Ala-Phe-7-amido-4-methylcoumarin (AAF-AMC) (Sigma #A3401) for 20 h at 37 °C. The reaction was stopped by 100 µl 0.5 M EDTA (pH 12.0). Fluorescence was measured using a BioTek Cytation 5 Cell Imaging Multi-Mode Reader with an extinction wavelength of 355 nm and an emission wavelength of 460 nm. To make a standard curve, the fluorescence intensity of different amounts of AMC was measured. For experimental samples, the fluorescence intensity was measured for each sample, and the enzyme activity was calculated as nmol/mg/h using the standard curve.

### 2.5. Statistical analysis

Unpaired two-tailed Student’s *t* tests were performed for all group comparisons using GraphPad Prism 6 software. Data are presented as the mean ± standard error of the mean (SEM). **** *p* < 0.0001.

## 3. Results

### 3.1. Generation of TPP1-mutant hESC lines using CRISPR/Cas9-based genome editing

The *TPP1* gene is located at the chromosome band 11p15.4, consisting of 13 exons and 12 introns and spanning 6.65 kb (Haines et al., 1998; Liu et al., 1998; Sharp et al., 1997). The two most frequently reported *TPP1* pathogenic variants that are associated with CLN2 disease and TPP1 enzyme deficiency are c.509-1G>C and c.622C>T (p.(Arg208*)) (Gardner et al., 2019; Sleat et al., 1999). The c.509-1G>C variant is a splice acceptor mutation that occurs in intron 5 of *TPP1*, whereas c.622C>T is a nonsense mutation that occurs in exon 6 (Fig. 1a). To expand the cellular models for investigating CLN2 disease, we introduced each of these two mutations separately in a founder hESC line (H9) using CRISPR/Cas9-mediated homology directed repair.

**Fig 1.**
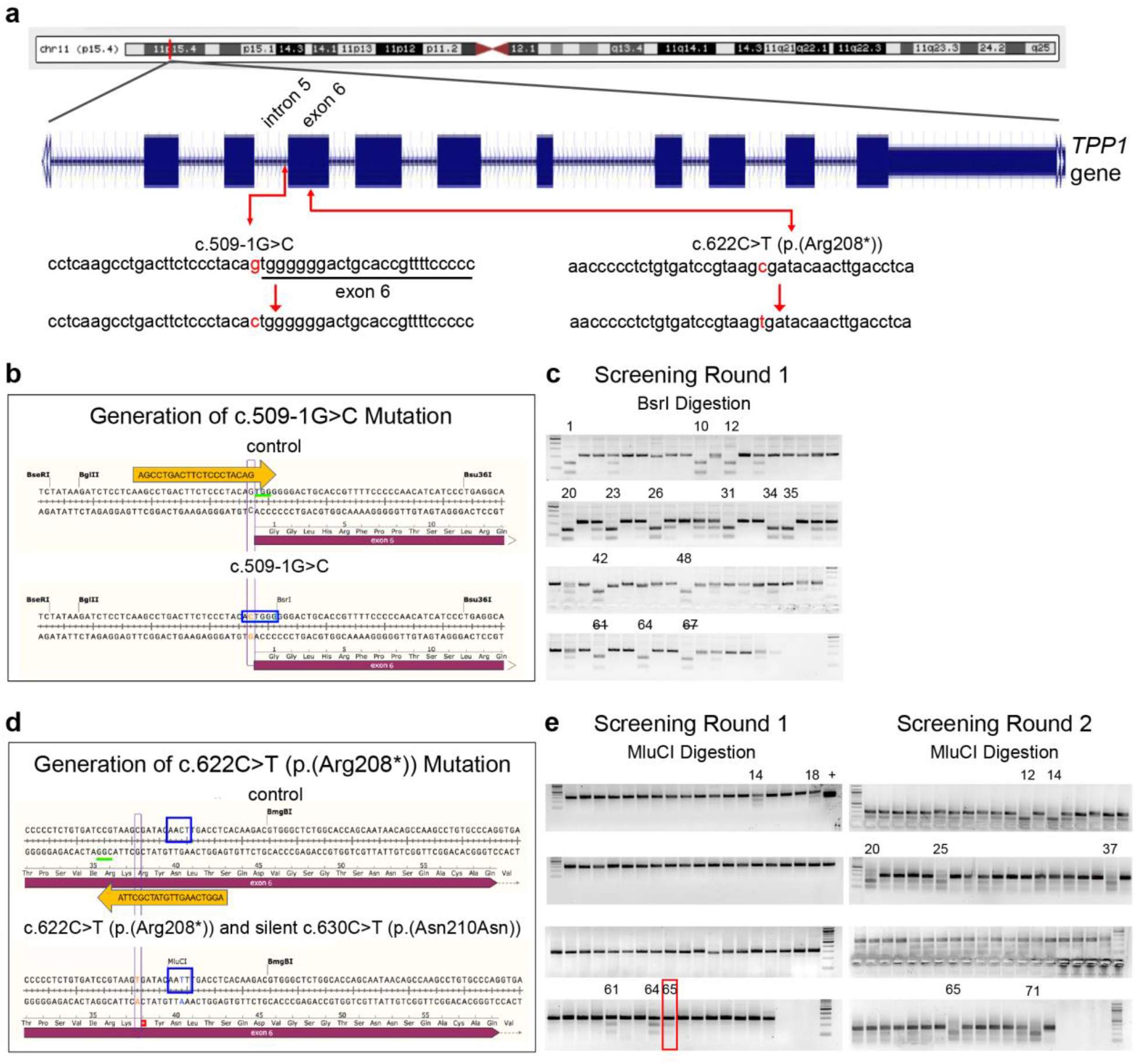
Generation of CLN2 disease-associated mutations in *TPP1* in H9 hESCs using CRISPR/Cas9 knock-in methodology. **(a)** Diagram of *TPP1* location on chromosome 11 [adapted from UCSC Genome Browser, GRCh38/hg38, http://genome.ucsc.edu (Kent et al., 2002)] and two common CLN2 disease-associated mutations in *TPP1*, the c.509-1G>C splice-site splice acceptor mutation and the c.622C>T (p.(Arg208*)) nonsense mutation. **(b, c)** Schematic of the CRISPR/Cas9 knock-in methodology used to generate the c.509-1G>C mutation in *TPP1* (b) and gels of PCR products digested using BsrI as a means for screening clones for successful targeting of alleles (c). **(d, e)** Schematic of the CRISPR/Cas9 knock-in methodology used to generate the c.622C>T (p.(Arg208*)) mutation in *TPP1* (d) and gels of PCR products digested using MluCI as a means for screening clones for successful targeting of alleles (e). Gels from the two rounds of screening are shown. Clone #65 (red box) from the first round of gene editing was identified as a heterozygous mutant and was retargeted in a second round of gene editing to generate homozygous mutant clones. Orange arrows depict gRNAs. Green underlines indicate the protospacer adjacent motif (PAM) sequences. Red font indicates the mutated nucleotides. Blue rectangles indicate the Bsr| restriction enzyme site created upon knock-in of the c.509-1G>C mutation (b) and the MluC| restriction enzyme site created upon knock-in of a silent mutation (to prevent re-cutting) (c.630C>T [p.(Asn210Asn)]) along with knock-in of the c.622C>T (p.(Arg208*)) mutation (d). Clone numbers are indicated above the gels. In validating clones identified by BsrI digestion to have the c.509-1G>C mutation, clone #61 was found to have an additional mutation upon sequencing and cells from clone #67 were phenotypically poor. Thus, these clone numbers are indicated with a strikethrough. The “+” above an MluCI digestion gel in the first round of screening indicates a positive control.

The gRNA and genome editing strategy for generation of the c.509-1G>C mutation are shown in Fig. 1b, and sequences are provided in Supplementary Table 1. This mutation created a BsrI (ACTGGG↓) restriction site. As such, complete digestion indicated that both alleles had been targeted (Fig. 1c). In the first round of screening, among 73 clones tested, 13 clones showed complete digestion and Sanger sequencing confirmed the desired editing of both alleles absent other mutations. That is, the clones were homozygous for the c.509-1G>C mutation, resulting in a homology directed repair efficiency of 17.8% (13/73). One of the 13 clones was found to consist of phenotypically poor cells, thus eventually, 12 clones were expanded and stocked. The c.509-1G>C mutation clones #1, #20, #23, and #42 were randomly selected for further studies (Fig. 1c).

The gRNA and genome editing strategy for generation of the c.622C>T (p.(Arg208*)) mutation are shown in Fig. 1d, and sequences are provided in Supplementary Table 1. In generating this mutation, a silent mutation was also inserted to prevent re-cutting of the genome (c.630C>T [p.(Asn210Asn)]). The silent mutation was designed so as to also create an MluC| (↓AATT) restriction site, which was used for clone screening (Fig. 1e). For this mutation, in the first round of screening, among 72 clones tested, 5 clones showed partial digestion. Sanger sequencing indicated that one allele was correctly targeted for the c.622C>T;630C>T mutations in these clones, whereas the other allele had an indel (4 clones), representing non-homologous end joining instead of homology directed repair, or was wild-type (1 clone). Two clones in which the second allele had an indel were selected for further analysis as compound heterozygous coding mutants (clones #14 and #64), and the clone in which the second allele was wild-type was selected as a heterozygous mutant for further analysis (clone #65) (Fig. 1e).

The heterozygous mutant from the first round of screening with one mutant allele and one wild-type allele (clone #65) was retargeted to generate homozygous mutant clones of the c.622C>T;630C>T mutations. In the second round of screening, among 72 clones tested, 7 showed complete (or nearly complete) digestion and Sanger sequencing confirmed the desired editing of both alleles absent other mutations, resulting in an overall homology directed repair efficiency of 4.9% (7/144). Two clones homozygous mutant for the c.622C>T;630C>T mutations were randomly selected for further analysis (clones #12 and #25).

### 3.2. Sanger sequencing of TPP1-mutant lines

Sanger sequencing chromatograms of PCR products covering the targeted regions of the *TPP1*-mutant lines are shown in Fig. 2. A chromatogram from a homozygous *TPP1* c.509-1G>C mutation line (clone #23) is shown in Fig. 2a. Chromatograms from a homozygous *TPP1* c.622C>T;630C>T (p.(Arg208*;Asn210Asn)) mutation line (clone #12) and the heterozygous *TPP1* mutation line c.[622C>T;630C>T];[622=;630=] (p.[(Arg208*;Asn210Asn)];[(Arg208=;Asn210=)]) (clone #65) are shown in Fig. 2b and c, respectively. Fig. 2d shows a chromatogram for the compound heterozygous coding mutation of the following genotype: c.[617_620delGTAA];[622C>T;630C>T] (p.[(Lys207Aspfs*210)];[(Arg208*;Asn210Asn)] (clone #14). Here, on one allele, deletion of GTAA from nucleotide 617 to 620 is predicted to cause a frameshift leading to missense changes from amino acid 207 to 209 and a nonsense mutation at position 210. On the other allele, the c.622C>T mutation leads to the p.(Arg208*) nonsense mutation and the c.630C>T mutation is the p.(Asn210Asn) silent mutation. Fig. 2e shows a chromatogram for the compound heterozygous coding mutation of the following genotype: c.[621_622insG];[622C>T;630C>T] (p.[(Arg208Alafs*223)];[(Arg208*;Asn210Asn)] (clone #64). On one allele, insertion of a G between nucleotide 621 and 622 is predicted to lead to a frameshift resulting in missense changes from amino acid 208 to 222 and a nonsense mutation at position 223. On the other allele, the c.622C>T mutation leads to the p.(Arg208*) nonsense mutation and the c.630C>T mutation is the p.(Asn210Asn) silent mutation. The locations of the mutations are based on TPP1 transcript NM_000391.4 and protein NP_ 000382.3. All references to genomic coordinates are based on human genome build hg38 (www.genome.ucsc.edu).

**Fig 2.**
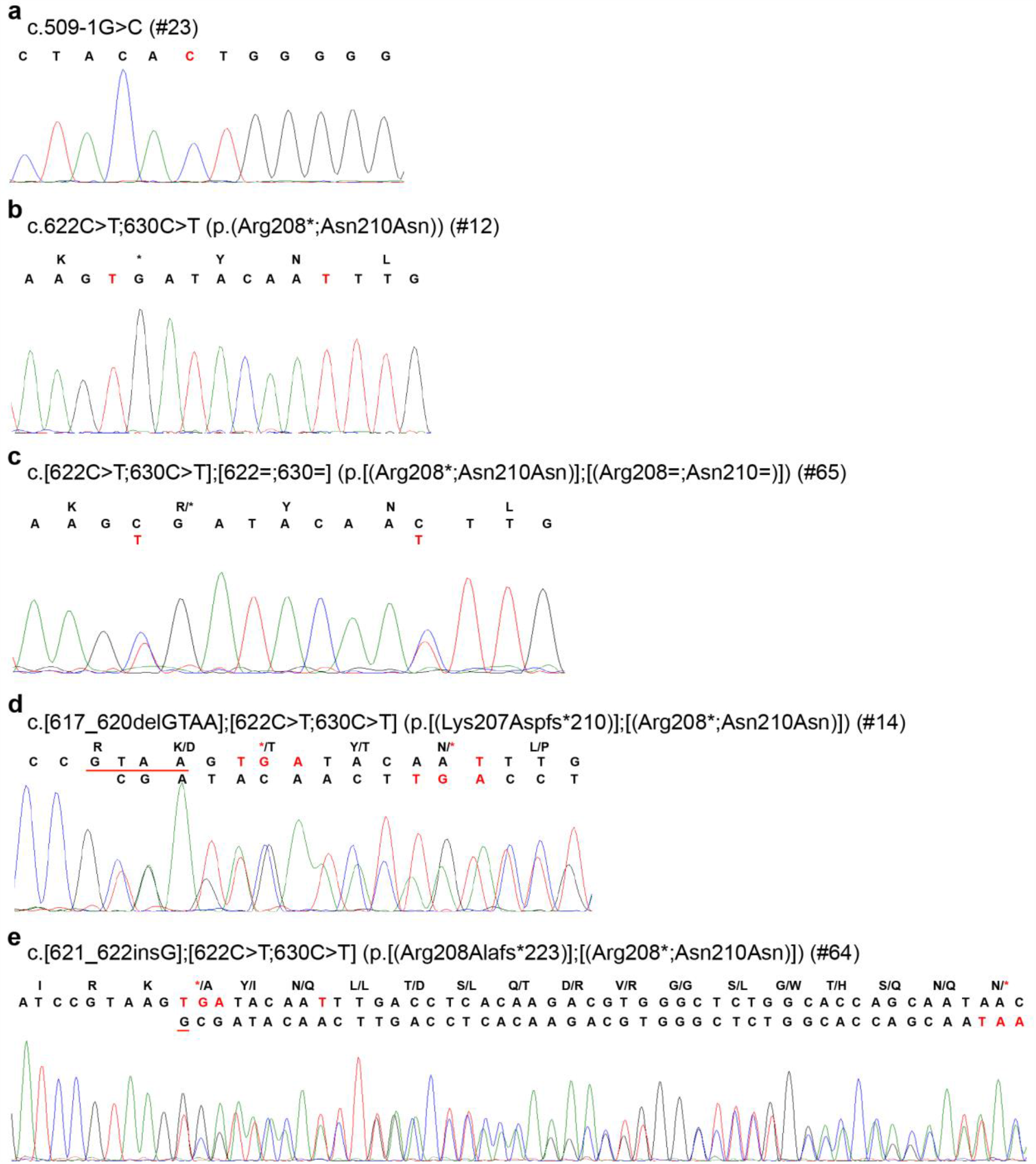
Sanger sequencing chromatograms of homozygous and heterozygous mutations induced in *TPP1* using CRISPR/Cas9 knock-in methodology. **(a-e)** Sanger sequencing chromatograms of *TPP1*-mutant hESC lines of the following genotypes, respectively: c.509-1G>C, c.622C>T;630C>T (p.(Arg208*;Asn210Asn)), c.[622C>T;630C>T];[622=;630=] (p.[(Arg208*;Asn210Asn)];[(Arg208=;Asn210=)]), c.[617_620delGTAA];[622C>T;630C>T] (p.[(Lys207Aspfs*210)];[(Arg208*;Asn210Asn)]), and c.[621_622insG];[622C>T;630C>T] (p.[(Arg208Alafs*223)];[(Arg208*;Asn210Asn)]). The clone numbers of the *TPP1*-mutant hESCs from which the chromatograms were derived are indicated. Red font indicates mutated nucleotides, and the deletion (d) and insertion (e) are underlined in red.

### 3.3. Stem cell morphology in the TPP1-mutant lines is comparable to the control line

The control H9 hESC line and the lines with different *TPP1* mutations were expanded and cultured in serum-free, feeder-free media and grown on Matrigel pre-coated plates. Representative bright field images of hESC colonies are shown in Fig. 3a-f. Control (Fig. 3a) and *TPP1*-mutant (Fig. 3b-f) hESCs grew as colonies consisting of cells tightly packed with each other. Also, the hESCs in the colonies comprising *TPP1*-mutant cells displayed classic pluripotent stem cell morphology, with a high nucleus to cytoplasm ratio, similar to hESCs in the control H9 colony.

**Fig 3.**
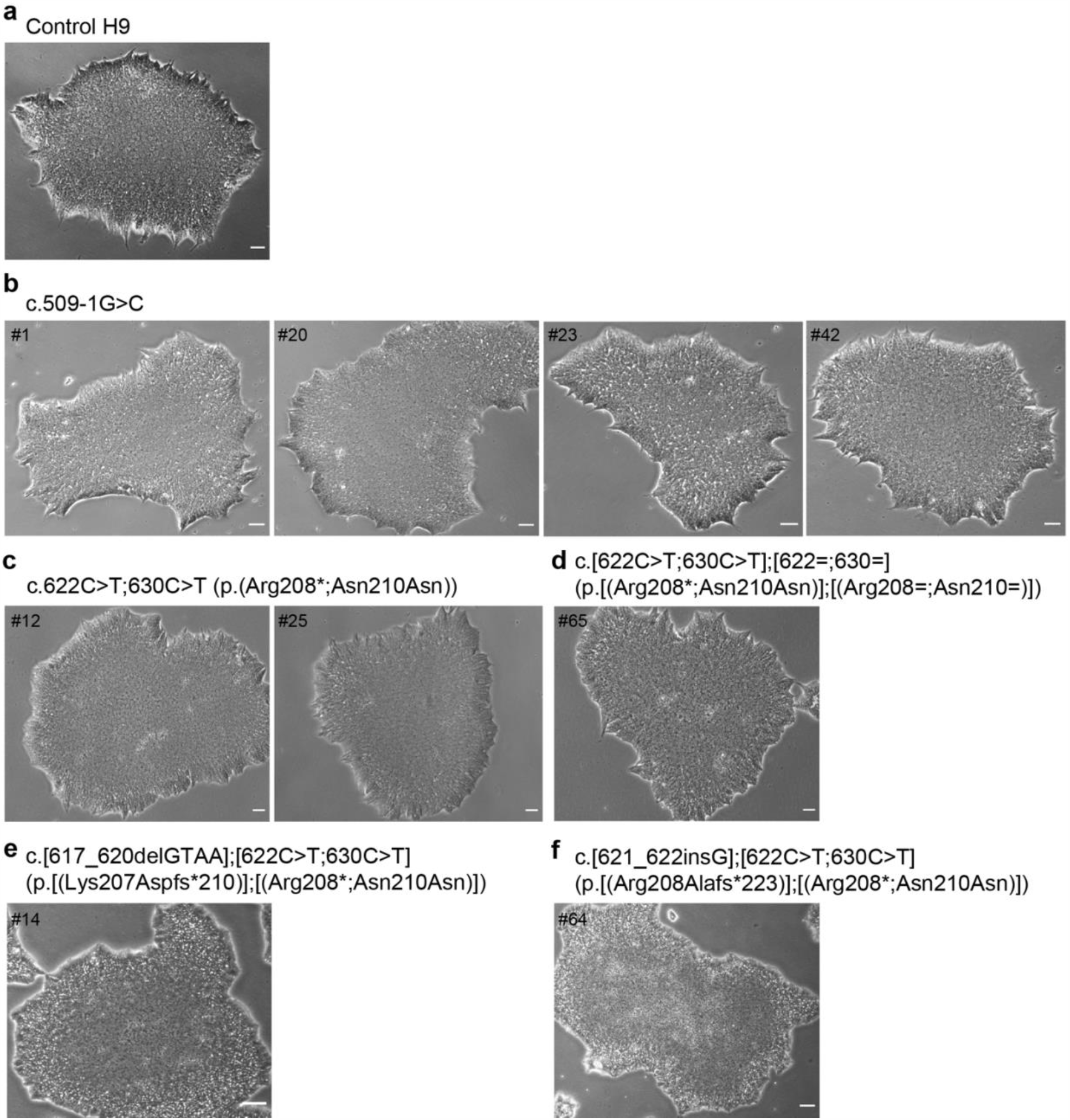
Morphology of cell colonies from the control H9 hESC line and *TPP1*-mutant hESC lines. **(a-f)** Bright field images of colonies of hESCs from the control H9 hESC line (a) and from *TPP1*-mutant hESC lines of the following genotypes: c.509-1G>C (b), c.622C>T;630C>T (p.(Arg208*;Asn210Asn)) (c), c.[622C>T;630C>T];[622=;630=] (p.[(Arg208*;Asn210Asn)];[(Arg208=;Asn210=)]) (d), c.[617_620delGTAA];[622C>T;630C>T] (p.[(Lys207Aspfs*210)];[(Arg208*;Asn210Asn)]) (e), and c.[621_622insG];[622C>T;630C>T] (p.[(Arg208Alafs*223)];[(Arg208*;Asn210Asn)]) (f). The clone numbers of the *TPP1*-mutant hESCs for which images of colonies are shown are indicated. Scale bar, 40 µM.

### 3.3. TPP1 mutations lead to absence of TPP1 protein and deficiency in TPP1 enzyme activity

The *TPP1* gene encodes the soluble lysosomal peptidase TPP1. Human TPP1 consists of 563 amino acid residues and contains three major domains, a signal peptide (amino acids 1 to 19), a propeptide (amino acids 20 to 195), and a peptidase domain (amino acids 196 to 563). The 19-amino acid signal peptide is cleaved off when the protein is translocated into the lumen of the endoplasmic reticulum, and a 176-amino acid propeptide is removed during the maturation process to yield a mature enzyme of 368 amino acid residues (Lin et al., 2001; Sleat et al., 1997). TPP1 contains five N-glycosylation sites inside the peptidase domain, which are necessary for appropriate folding, protein processing, and enzymatic activity (Golabek et al., 2004; Lin et al., 2001; Pal et al., 2009; Sleat et al., 1997; Wujek et al., 2004).

Given the association of pathogenic *TPP1* mutations with CLN2 disease, we analyzed TPP1 protein expression and enzyme activity in our *TPP1*-mutant hESC lines. For protein expression studies, cells were harvested and lysates were extracted from control cells (H9 hESCs and HEK293 cells) and from *TPP1*-mutant hESCs. Protein amounts and molecular weights were then determined by Western blot using an antibody against amino acids 195 to 305 of human TPP1, which should thus recognize the peptidase domain.

As shown in Fig. 4a, in control cells, the proenzyme form of TPP1 was detected by Western blot at a molecular weight of ∼71 kDa (top red arrow, middle panel), and the processed mature form was detected at a molecular weight of ∼48 kDa (bottom red arrow, middle panel). Similarly, in heterozygous mutant c.622C>T;630C>T (p.(Arg208*;Asn210Asn)) hESCs, both the proenzyme and mature forms of TPP1 were found to be present (red arrows, left panel), albeit in reduced amounts. To this end, in comparison to control H9 hESCs, the following amounts of TPP1 were detected in lysates from heterozygous mutant hESCs: 27.7% (proenzyme), 36.1% (mature), and 33.6% (proenzyme + mature). In contrast, TPP1 was no longer detected, in either the proenzyme or mature form, in homozygous mutant c.622C>T;630C>T (p.(Arg208*;Asn210Asn)) hESCs. Regarding lysates from hESCs containing the homozygous *TPP1* c.509-1G>C splice acceptor mutation, we observed two bands, neither of which directly corresponded to the two bands present in control H9 hESCs. The upper band ran at a molecular weight just below that of the control proenzyme form (top green arrow, middle panel), whereas the lower band ran at a molecular weight above that of the control active form (∼60 kDa, bottom green arrow, middle panel).

**Fig 4.**
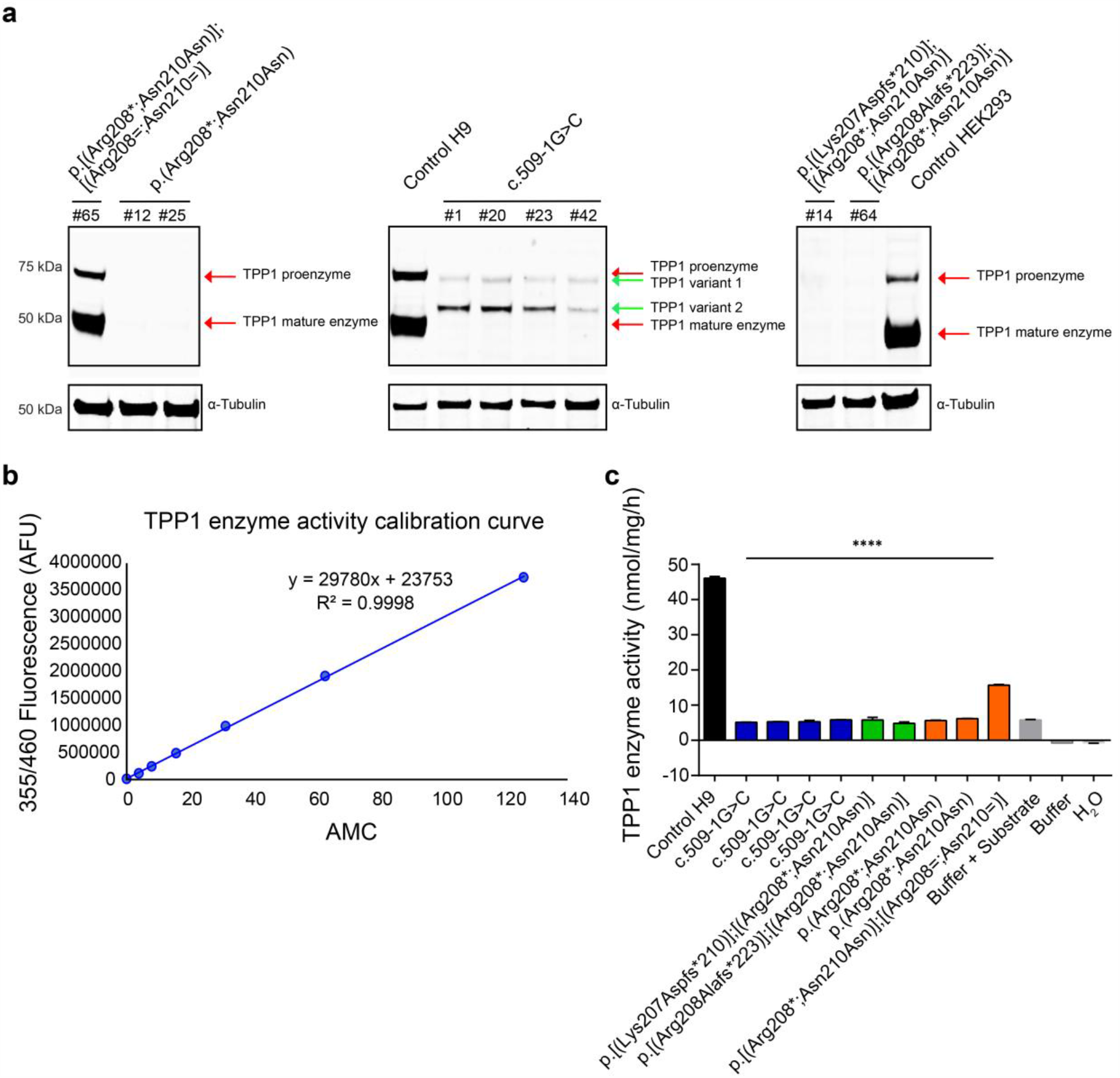
Analysis of protein expression and TPP1 enzyme activity in control H9 hESCs and *TPP1*-mutant hESCs. **(a)** Western blot of lysates from control cells (H9 hESCs and HEK293 cells) and *TPP1*-mutant hESCs of the indicated genotypes using antibodies against TPP1 and α-Tubulin (as a loading control). Lysates from HEK293 cells were used as a non-stem cell line control. Note, the Western blot was run as a single gel; however, it has been separated into three sections so as to be able to more clearly indicate the different TPP1 bands (red and green arrows). **(b)** Calibration curve used for calculating TPP1 enzyme activity. The curve was generated by determining the fluorescence intensity of emission at 460 nm, graphed as arbitrary fluorescence units (AFU), for different amounts of AMC (0, 3.9, 7.8, 15.625, 31.25, 62.5 and 125 nmol). **(c)** TPP1 enzyme activity in lysates from control H9 hESCs and from *TPP1*-mutant hESCs of the indicated genotypes, as determined by AMC production. Buffer with or without substrate and H_2_O were used as controls. Lysates from cells or buffer were incubated with AAF-AMC at 37 °C for 20 h; enzyme activity was normalized to total protein and is represented as nmol/mg/h. *n* = 3 replicates. Data are presented as means ± SEMs. Unpaired Student’s *t* tests were used. **** *p* < 0.0001.

AAF-AMC (Ala-Ala-Phe-7-amido-4-methylcoumarin) is a fluorogenic aromatic tripeptide substrate that can be cleaved by serine proteases and used to assay enzyme activity. Namely, once cleaved, AAF-AMC generates the fluorescent chromophore AMC (7-amino-4-methylcoumarin). TPP1 enzyme activity was measured based in use of AAF-AMC as a substrate and according to the fluorescence intensity of AMC (Fig. 4b, c). The amount of product cleaved by cell lysates derived from the control H9 hESC line and from the *TPP1*-mutant lines was calculated based on the equation derived from the standard curve (Fig. 4b), and TPP1 enzyme activities were graphed as nmol product cleaved/mg protein/h (Fig. 4c). Cell lysates from the control H9 hESC line showed the highest amount of enzyme activity (46.06 ± 0.54 nmol/mg/h), as expected. Lysates from heterozygous mutant c.622C>T;630C>T (p.(Arg208*;Asn210Asn)) hESCs showed TPP1 enzyme activity, but at a decreased rate (15.69 ± 0.17 nmol/mg/h) in comparison to control H9 hESCs. In contrast, lysates from all other *TPP1*-mutant hESCs showed a rate of enzyme activity below or close to 5 nmol/mg/h, comparable to the buffer + substrate control (Fig. 4c).

## 4. Discussion

We report here on two high-incidence CLN2 disease-associated *TPP1* mutations we generated in a founder hESC line (H9), using CRISPR/Cas9 homology directed repair knock-in technology, a c.509-1G>C splice acceptor mutation and a c.622C>T (p.(Arg208*)) nonsense mutation. In generating these mutations in a homozygous state, the c.622C>T (p.(Arg208*)) mutation was also generated in the heterozygous state (c.[622C>T;630C>T];[622=;630=] (p.[(Arg208*;Asn210Asn)];[(Arg208=;Asn210=)])). Additionally, two coding mutations were generated in a compound heterozygous state as a result of one allele undergoing non-homologous end joining; these mutations were: c.[617_620delGTAA];[622C>T;630C>T] (p.[(Lys207Aspfs*210)];[(Arg208*;Asn210Asn)]) and [621_622insG];[622C>T;630C>T] (p.[(Arg208Alafs*223)];[(Arg208*;Asn210Asn)]). Mutations were confirmed by Sanger sequencing, and colonies of control H9 hESCs vs. *TPP1*-mutant hESCs were of similar morphology. At the protein level, heterozygous mutant c.622C>T;630C>T (p.(Arg208*;Asn210Asn)) hESCs expressed proenzyme and mature forms of TPP1 in a manner similar to control H9 hESCs, whereas in the homozygous state, no TPP1 was detected. TPP1 protein was also no longer detected in hESCs containing the compound heterozygous coding mutations. In contrast, c.509-1G>C hESCs expressed TPP1 in forms that were detected as two bands on Western blotting, neither of which clearly corresponded to the proenzyme or mature forms of TPP1 present in control H9 hESCs. TPP1 enzyme activity was measured in vitro using lysates from hESCs. Heterozygous mutant c.622C>T;630C>T (p.(Arg208*;Asn210Asn)) hESCs showed reduced enzyme activity, whereas all other *TPP1*-mutant hESCs examined failed to exhibit any TPP1 enzyme activity above background. Thus, *TPP1* mutations in these lines represent loss-of-function mutations.

Mutations occurring in nucleotides surrounding a 5’ or 3’ splice site can lead to mis-splicing and typically result in exon skipping, activation of a cryptic splice site, or intron retention (Anna and Monika, 2018; Epstein et al., 1993; Ward and Cooper, 2010; Wimmer et al., 2007). Most of the splice site mutations that lead to human disease involve the invariant intronic GT and AG dinucleotides in the 5’ and 3’ splice sites (Krawczak et al., 2007; Ramalho et al., 2003; Symoens et al., 2011). Regarding TPP1 detected in lysates from c.509-1G>C hESCs, that is, cells with the *TPP1* splice acceptor mutation, it is possible that the two observed bands reflect protein products resulting from splicing abnormalities arising due to the mutation. Further studies are needed, however, to examine the splicing mechanism in mutant cells and to determine the identity of these bands.

Aberrant accumulation of mitochondrial ATP synthase subunit C in lysosomes is a feature of CLN2 disease (Ezaki et al., 1996; Palmer et al., 1992; Palmer et al., 1989), as well as other NCLs. For example, NCLs caused by *CLN6* mutation and *CLN3* mutation share clinical and pathological features, including lysosomal accumulation of ATP synthase subunit C in Lamp 1-positive organelles from cerebellum cells (Cao et al., 2011). Indeed, the pathological hallmark of NCL caused by *CLN3* mutation is autofluorescent ceroid lipofuscin deposits within autolysosomes that are enriched in subunit C of the mitochondrial ATP synthase complex (Fossale et al., 2004; Kominami, 2002; Palmer et al., 1992). These cellular features of mutant neurons will be examined in future studies.

For our studies we used a normal, female hESC line (H9) as the cellular context into which *TPP1* mutations were induced. Patient-derived iPSCs also provide highly useful, biologically relevant models for study of neurological disorders and development of therapeutics (Young-Pearse and Morrow, 2016). Indeed, two prior studies have reprogrammed fibroblasts from patients with CLN2 disease to iPSCs, which were then differentiated to neuronal progenitor cells and mature neurons or neuronal stem cells as a means for disease modeling and testing of potential therapeutics (Lojewski et al., 2014; Sima et al., 2018). In characterizing disease pathology, these studies found that neuronal precursors and mature neurons in particular displayed NCL-like characteristics with respect to organellar pathology (i.e., aberrant endosomes/lysosomes) and accumulation of lipids and autofluorescent storage material. These patient-derived, cellular CLN2 disease models were also used as platforms to test potential therapies – small molecules and TPP1 protein replacement. The studies by Lojewski et al. (2014) and Sima et al. (2018) support that patient-derived neurons, by way of iPSCs, are of great benefit as a human cellular model for the study of CLN2 disease pathology and for genotype-driven drug development. At the same time, they present the limitations that, in each case, compound heterozygous mutations were modeled and the models did not have associated isogenic control lines. As such, study of a single specific CLN2 disease mutation and comparison of results to a direct genetically related control cannot be performed using these patient-derived iPSC lines. Furthermore, the parent cell line used in the studies presented here is a hESC line, as compared to iPSCs generated from mature fibroblasts in these prior studies. The H9 hESC line is a very commonly used line and robustly tested for studies of neuronal development and other stem cell applications (Erceg et al., 2008; Liu et al., 2015; Marei et al., 2017).

In summary, we induced *TPP1* mutations, including the two most commonly associated with CLN2 disease, in the H9 hESC line using CRISPR/Cas9 technology and analyzed protein expression and enzyme activity in the generated cell lines. The results are supportive of generation of loss-of-function *TPP1*-mutant hESCs. Thus, these hESC models will be useful for future studies of molecular and cellular mechanisms underlying CLN2 disease and for therapeutic development. To this end, we are making our c.509-1G>C and c.622C>T (p.(Arg208*)) *TPP1*-mutant hESC lines available for wide-spread sharing by depositing them with the WiCell biorepository.

## Author contributions

Li Ma: Formal analysis, Investigation, Writing – Original Draft, Visualization. Adriana Prada: Investigation, Writing – Original Draft, Visualization. Michael Schmidt: Investigation, Data curation. Eric M. Morrow: Conceptualization, Methodology, Formal analysis, Writing – Review & Editing, Supervision, Funding acquisition.

## Acknowledgements

The authors acknowledge the support of the Sandra Nusinoff Lehrman and Stephen Lehrman Family Fund provided to the Morrow Laboratory at Brown University that led to the development of the *TPP1*-mutant stem cell lines. The sponsor had no role in study design; in the collection, analysis and interpretation of data; in the writing of the report; or in the decision to submit the article for publication. The authors also acknowledge Heather M. Thompson, Ph.D., of Brown University for assistance in manuscript preparation and assistance from the Stem Cell Core at the University of Connecticut Health Center.

## Supplementary data

Supplementary Table 1. gRNA and ssODN sequences. Nucleotides to be edited in generating the c.509-1G>C and c.622C>T mutations are indicated in red font. The silent c.630C>T mutation in the ssODN sequence used for generating the c.622C>T mutation is indicated in blue font.

Supplementary Table 2. PCR primer sequences.

Supplementary Table 3. List of *TPP1*-mutant hESC lines and clone numbers.

## Footnotes

1 AAF-AMC: Ala-Ala-Phe-7-amido-4-methylcoumarin; AFU: Arbitrary fluorescence units; AMC: 7-amino-4-methylcoumarin; CLN: Ceroid lipofuscinosis, neuronal; CRISPR: Clustered regularly interspaced short palindromic repeats; Cas9: CRISPR-associated protein 9; gRNA: Guide RNA; hESC: Human embryonic stem cell; iPSC: Induced pluripotent stem cell; NCL: Neuronal ceroid lipofuscinosis; PAM: Protospacer adjacent motif; ROCKi: Rho-associated protein kinase inhibitor; ssODN: Single-stranded oligodeoxyribonucleotide; TPP1: Tripeptidyl peptidase 1.

